## Supplementary material for "Real-time feeding behavior monitoring by electrical penetration graph rapidly reveals host plant susceptibility to crapemyrtle bark scale (Hemiptera: Eriococcidae)": Figure S1: EPGminer application released on Zenodo and GitHub assists in semi-automatical calculation of feeding characteris-tics and visualization of

### Supplementary Materials

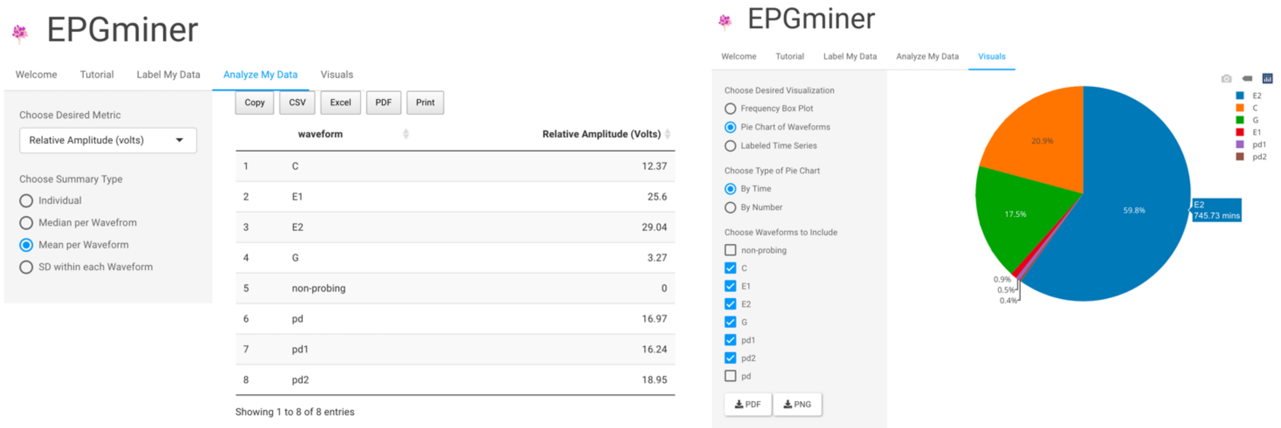


**Figure S1.** EPGminer application released on Zenodo and GitHub assists in semi-automatical calculation of feeding characteristics and visualization of EPG analysis results for feeding behavior study of *Acanthococcus lagerstroemiae.* EPGminer has the website version https://epgdata.shinyapps.io/epgminer_app/ and the software version. R package code is available in Zenodo: https://doi.org/10.5281/zenodo.6299917. https://github.com/LylChun/epgminer/tree/master/inst/epgminer_app/rsconnect/shinyapps.io/epgdata. Detailed instructions on EPGminer utilization (step-by-step demo included) in the calculation of frequency and relative amplitude for each waveform are provided in GitHub https://lylchun.github.io/EPGminer/index.html.

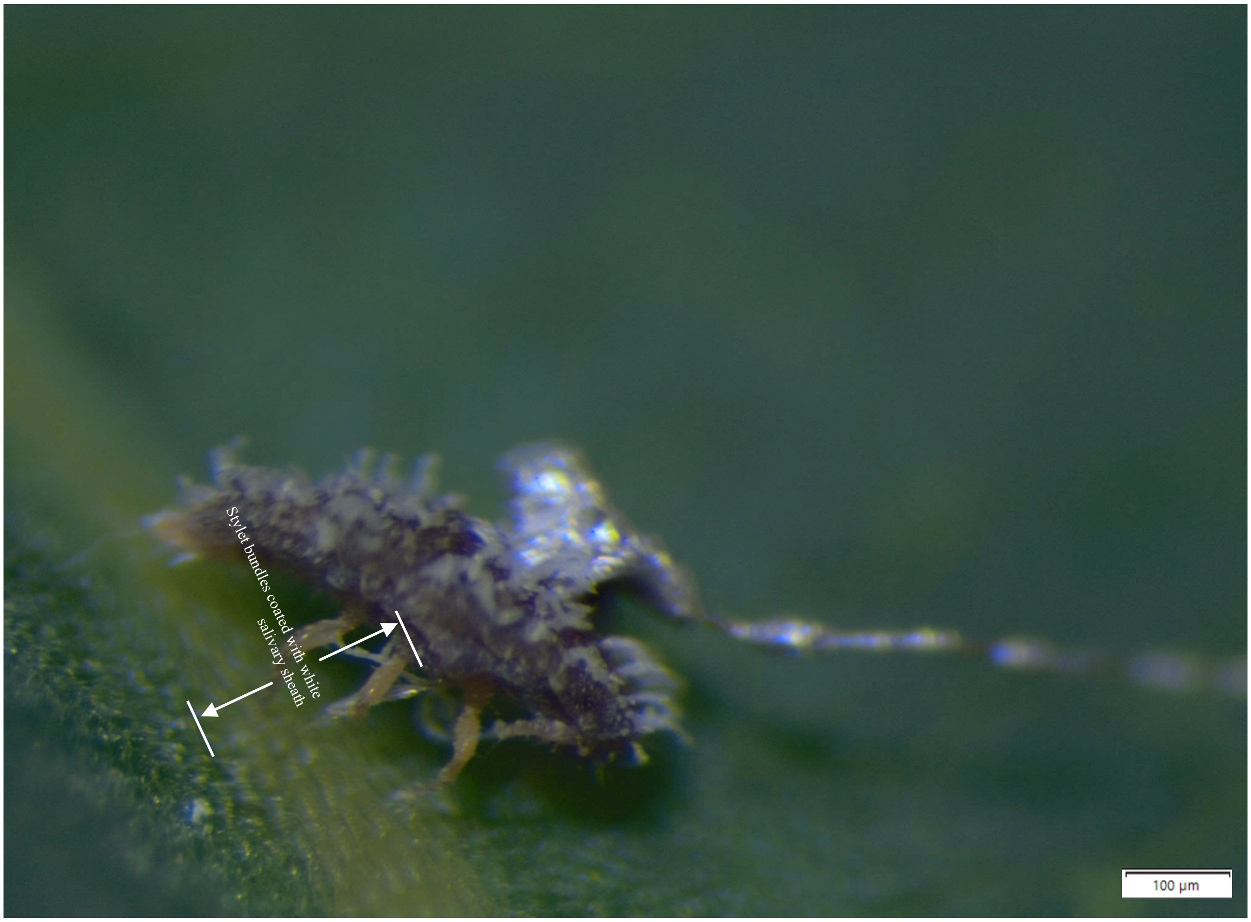


**Figure S2.** *Acanthococcus lagerstroemiae* penetrates its stylet bundles coated with salivary sheath into a plant. It was observed that *A. lagerstroemiae* penetrated its stylet bundles coated with white salivary sheath when feeding on a plant (photographed under an Olympus SZX7 stereomicroscope coupled with an Olympus LC30 camera (Waltham MA, United States)).

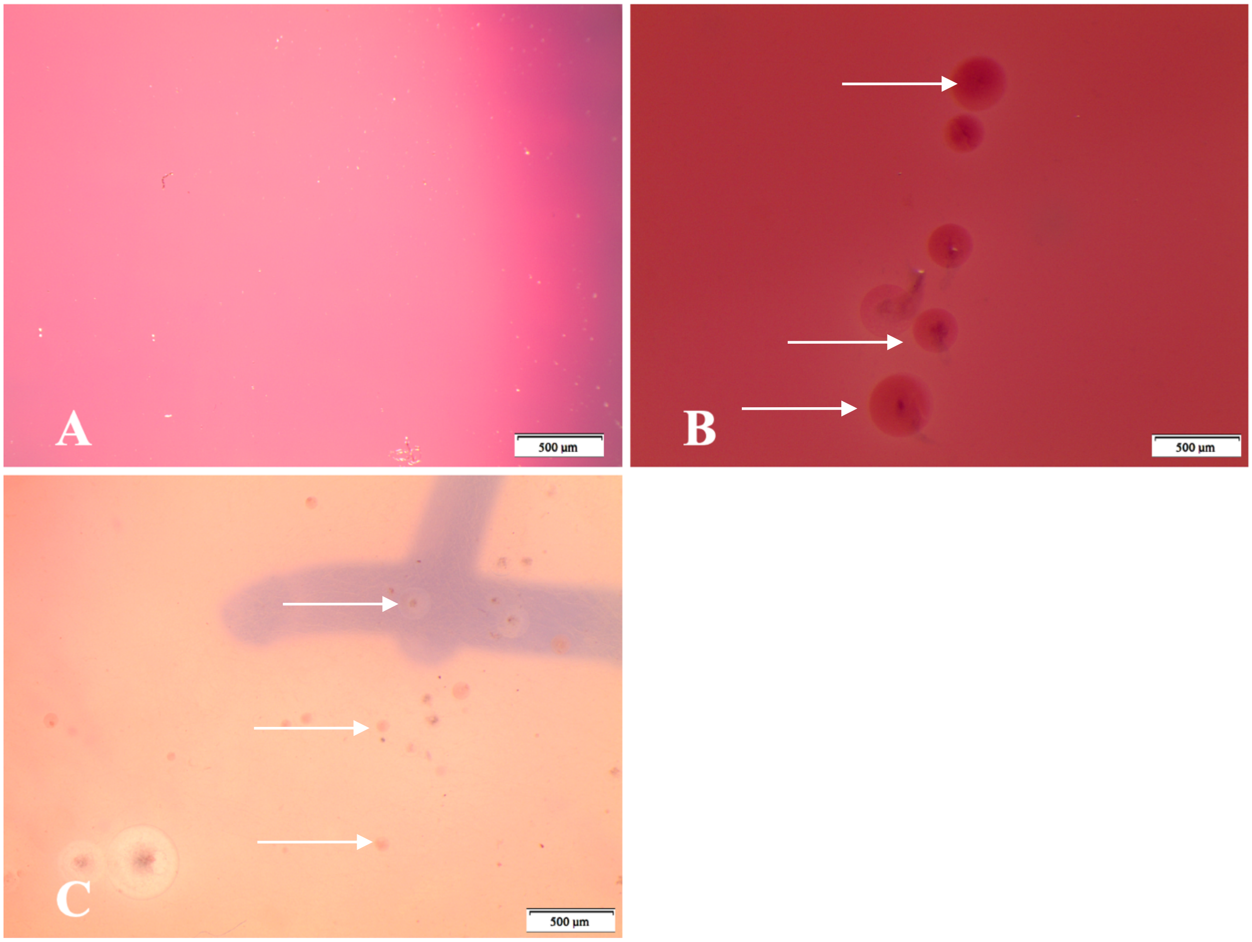


**Figure S3.** Overview of pectinesterase detection in agarose gel plates. (**A**) No rings or halos were shown when *A. lagerstroemiae* was feeding on the plates with no pectin added. (**B**) Maroon-stained rings and halos (arrows), representing pectinesterase activity, were shown by using ruthenium red around the feeding site of the stylet sheath when *A. lagerstroemiae* was feeding on 0.1% pectin-added gel plates. (**C**) The number and size of rings and halos in 1.0% pectin-added plates were not different from 0.1% pectin plates.
